# Evolution toward larger cell size reduces internal cellular density

**DOI:** 10.1101/2025.11.05.686806

**Authors:** A. Sastokas, W.-C. Ho, K. Schmidlin, M. Bhatt, P. Brown, P. Crossland, A. Kindelin, Z. Uzum, D. Shepherd, M. Lynch, K. Geiler-Samerotte

## Abstract

Cellular density depends on the balance between cell volume and the amount of material contained within that volume. Cells generally maintain this balance during growth, across the cell cycle, and in response to environmental changes. But is density also maintained when cell size changes through evolution? We selected barcoded populations of *Saccharomyces cerevisiae* for increased cell size over approximately 160 generations and asked whether cellular material increased proportionally to preserve density. Evolved lineages became larger through multiple genetic routes, yet repeatedly converged on the same physiological outcome: larger evolved cells had lower internal density. Optical diffraction tomography showed that internal density declined as cell size increased, indicating that cellular material did not increase proportionally with cell size. Lower density was also observed in clonally propagated evolved isolates, indicating that it was a stable, heritable feature rather than a temporary response to selection. We next asked whether the same relationship extends over longer evolutionary timescales. Across published dry-mass and volume measurements from 68 species, larger cells contained more total material, but material again increased more slowly than volume. Consequently, cellular density declined with increasing cell size across species, paralleling the pattern observed during short-term experimental evolution. In sum, these results show that early adaptation to larger size can repeatedly uncouple volume expansion from material accumulation. Although cells found multiple genetic routes to become larger, volume repeatedly increased faster than cellular material, and that same scaling relationship is evident across species. Reduced density may therefore be a common consequence of evolving larger cell size.

## Introduction

Increasing cell size creates a basic scaling problem: the added volume must be filled with cellular material. Budding yeast routinely span substantial size differences: mean cell volume varies approximately 2.5-fold across growth rates, diploid cells are typically 1.5–1.8-fold larger than comparable haploids, and total cell volume can approach a twofold increase between birth and division (Adams 1977; Tyson et al. 1979; Soifer et al. 2016). Yet mass and volume generally remain coordinated across these states (Godin et al. 2010; Odermatt et al. 2021; Liu et al. 2022; Rollin et al. 2023), and cellular density varies by only approximately 2% over the cell cycle (Bryan et al. 2010). This density homeostasis requires the production of proteins, nucleic acids, lipids, and other cellular material to remain coordinated with volume expansion, which also depends on water, ions, and small osmolytes (Rollin et al. 2023). This balance matters because intracellular concentration determines molecular crowding and thereby affects diffusion, signaling, and other biochemical processes; both compression and dilution can disrupt cellular function (Miermont et al. 2013; Neurohr and Amon 2020).

This balance can break down when cells are forced to grow beyond their usual size over a short timescale. In *cdc28*-arrested budding yeast, cell volume increased almost tenfold; at these sizes, RNA and protein production no longer scaled with volume, causing cytoplasmic dilution and functional defects (Neurohr et al. 2019; Neurohr and Amon 2020). In an isogenic series of haploid through tetraploid yeast, median cell volume increased 3.8-fold, whereas total protein content increased only 2.95-fold, revealing sublinear proteome scaling with ploidy (Yahya et al. 2022). These studies show that particular engineered routes to enlargement can disrupt molecular scaling. They do not reveal whether diverse, growth-compatible mutations favored during early adaptation to larger size repeatedly produce the same shift in whole-cell material density.

Indeed, recent experimental evolution in *Saccharomyces cerevisiae* showed that cells can evolve a fourfold decrease in volume while maintaining robust size homeostasis and relatively high fitness (Garoña et al. 2026). This result demonstrates that large changes in size do not necessarily disrupt cellular function. Whether cells can accommodate evolutionary change in the opposite direction remains unknown. Does selection for increased size preserve the usual relationship between cellular material and volume, or does density shift as size evolves? Answering this question would be strengthened by observing many independently evolving lineages rather than relying on a single engineered mutation, because different genetic routes to increased size may reorganize cells in different ways.

Here, we used a massively parallel experimental evolution system to select barcoded yeast populations for increased cell size. We followed changes in size, shape, growth, and final culture optical density over approximately 160 generations, and then used optical diffraction tomography to measure internal density in individual evolved cells. Selection reproducibly produced larger, more elongated cells that maintained rapid growth but reached lower final optical densities. At the single-cell level, larger cells had lower internal refractive indices, showing that cellular material did not increase in proportion to cell size. This lower-density state was not a temporary response to ongoing size selection. It persisted after populations were frozen and revived and was recovered in clonally propagated isolates from the final timepoint of the experiment. The evolved large-cell populations also showed stable, environment-dependent tradeoffs, including greater sensitivity to elevated temperature. We then asked whether the inverse relationship between size and density extended beyond evolved yeast. Across 68 species spanning diverse branches of the tree of life, dry mass increased more slowly than cell volume, so larger cells were less dense. Together, our results suggest that density is regulated during growth but not fixed over evolution. Selection produced a stable shift toward lower density, and the same relationship between size and density appears across species.

## Results

### Selection for increased cell size tends to remodel cell geometry

To determine how cellular architecture changes as cell size increases, we conducted a massively parallel evolution experiment using an ancestral population containing approximately 300,000 uniquely barcoded *Saccharomyces cerevisiae* lineages (Levy et al. 2015; Schmidlin et al. 2024). The DNA barcodes act as heritable lineage identifiers, allowing independently evolved lineages to be distinguished. From the same ancestral pool, we initiated four populations in 6-mL cultures of glucose-limited M3 medium: two subjected to Large Size Selection (LS) and two subjected to No Size Selection (NS).

Every 48 hours, one million cells from each population were isolated by flow cytometry and transferred into fresh medium. In the LS populations, only cells within the upper 10% of the forward-scatter-area (FSC-A) distribution were eligible for transfer. Because FSC-A increases with cell size (Zakhartsev and Reuss 2018), this procedure repeatedly enriched the LS populations for larger cells. In the NS populations, one million cells were selected without regard to FSC-A. The NS populations therefore controlled for repeated growth, flow-cytometric sorting, and population bottlenecks without directly selecting on size. This procedure was repeated for 20 transfers over 38 days, corresponding to approximately 160 generations (**Fig. 1A**).

**Figure 1.**
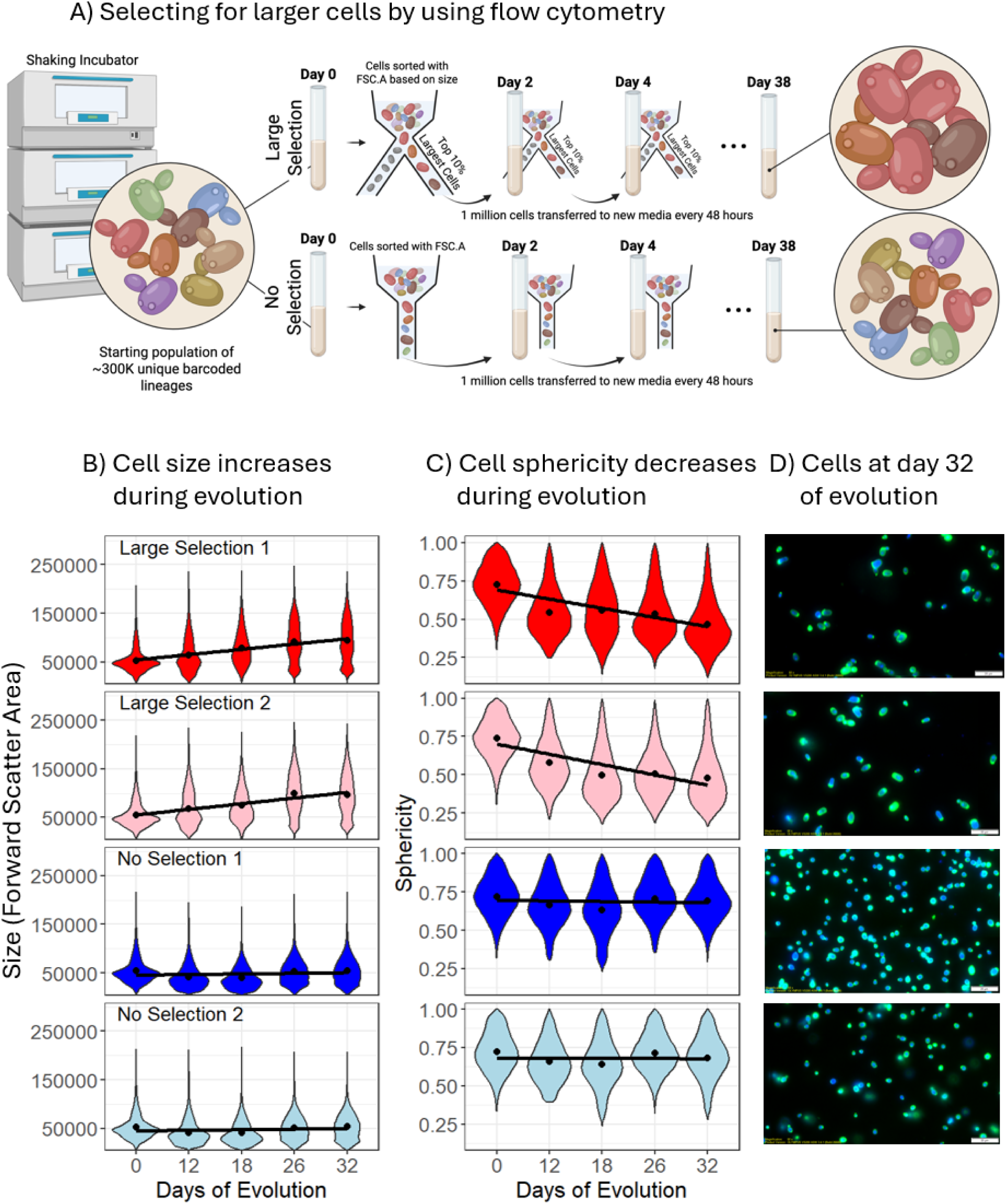
Experimental evolution generated changes in cellular size. A) A method cartoon depicting the design of the ‘large size selection’ and ‘no size selection’ evolution experiments. Two replicates were performed per selection regime. Each replicate started with 6mL of growth media and 1 million yeast cells from a pool that contained ∼300,000 unique barcodes. These cultures were incubated for 48 hours before being passed through a flow cytometer to measure cell size (forward scatter area, FSC-A). In the large size selection replicates, 1 million large cells with forward scatter measurements in the top 10% of the population were transferred to the next incubation vessel. In the no size selection replicates, 1 million random cells were transferred. This process continued for a total of 38 days. B) Violin plots of cell size across the large and no selection evolutions, including both replicates, at days 0, 12, 18, 26, and 32, with a linear regression of the mean values. C) Violin plots of cell sphericity across the large and no selection evolutions, including replicates, at days 0, 12, 18, 26, and 32, with a linear regression of the mean values. D) Microscopy images taken at day 32 of the evolution experiment for large and no selection evolutions, including replicates.

FSC-A increased progressively in both LS populations, whereas neither NS population showed a comparable trend (**Fig. 1B**). Regressions of mean FSC-A against evolutionary time had positive slopes in both LS populations and slopes near zero in the NS populations. By the final time point, mean FSC-A had increased by approximately 2-fold in LS1 and LS2 relative to the ancestor (**Fig. 1B**). Because forward scatter also depends on cell shape, these values quantify the response to selection but should not be interpreted as equivalent fold changes in cell volume. Still, repeated selection on FSC-A produced a reproducible increase in apparent cell size that was not caused simply by repeated growth and sorting.

Becoming larger creates a basic problem for a cell: proteins, nucleic acids, membranes, and other cellular components must increase along with cell volume if their intracellular concentrations are to remain approximately constant. Cells normally scale transcription and other biosynthetic processes with cell size (Sun et al. 2020; Cadart and Heald 2022; Swaffer et al. 2023). When this scaling breaks down in oversized cells, cytoplasmic dilution can impair cellular function (Neurohr et al. 2019; Neurohr and Amon 2020). Cell geometry may help shape this scaling challenge because increases in size need not occur uniformly in all directions. Across diverse cell types, elongation is a common growth strategy in bacteria, fungi, and other walled cells (Chan and Marshall 2010; Julien and Boudaoud 2018; Bozdag et al. 2023). In budding yeast, prolonged polarized growth can generate large, elongated cells, and diploid cells are both larger and more elongated than comparable haploid cells (Adams 1977; Ahn et al. 1999; Mable 2001; Klis et al. 2014). However, increased size does not require elongation: yeast cells can also enlarge while remaining relatively spherical, and aging polyploid cells can become more spherical as they continue to increase in size (Zadrag-Tecza et al. 2018). We therefore asked how cell geometry changed during selection for increased size.

We imaged population samples collected on days 0, 12, 18, 26, and 32 and quantified cell sphericity as the ratio of the minor to major cellular radius. A value of one represents a circular cell profile, whereas lower values indicate greater elongation. Mean sphericity declined progressively in both LS populations but remained comparatively stable in the NS populations (**Fig. 1C**). Microscopy likewise showed a reproducible population-level shift toward larger, more elongated cells in both LS populations by day 32, whereas the NS populations retained an average morphology more similar to that of the ancestor (**Fig. 1D**).

Cells under selection for increased size therefore did not simply expand proportionally in all dimensions. Instead, both LS populations showed a repeatable average shift toward elongation. By increasing one dimension more than others, elongation can increase cell size in a way that limits the corresponding increase in internal volume. This geometry could therefore reduce the amount of additional cellular material needed to maintain internal density as cells become larger.

### Larger cells preserve maximum growth rate but reach lower optical density

Recent experimental evolution of miniature yeast showed that cells can establish a dramatically smaller size while maintaining size homeostasis and relatively high competitive fitness (Garoña et al. 2026). That study demonstrated that substantial evolutionary changes in size do not necessarily cause a general loss of cellular function. We therefore first asked a complementary question: could cells selected for increased size also preserve basic growth performance?

Throughout the evolution experiment, we measured optical density at 600nm (OD_600_) every 10 min during each 48-h growth cycle (**Fig. 2A**). Consistent with typical microbial growth, the OD slowly increased at the beginning of each cycle before entering a phase of fast growth until saturation. Maximum growth rate per cycle increased over evolutionary time in both the LS and NS populations (**Fig. 2B**), consistent with the frequent evolution of faster growth during repeated propagation under a fixed laboratory condition (Jasmin and Zeyl 2012; Sezmis et al. 2018; Garoña et al. 2026). The parallel response under both selection regimes indicates that faster growth was a general adaptation to the experimental environment rather than an effect unique to increased cell size. Importantly, the LS populations were able to evolve larger size without losing their ability to grow rapidly.

**Figure 2.**
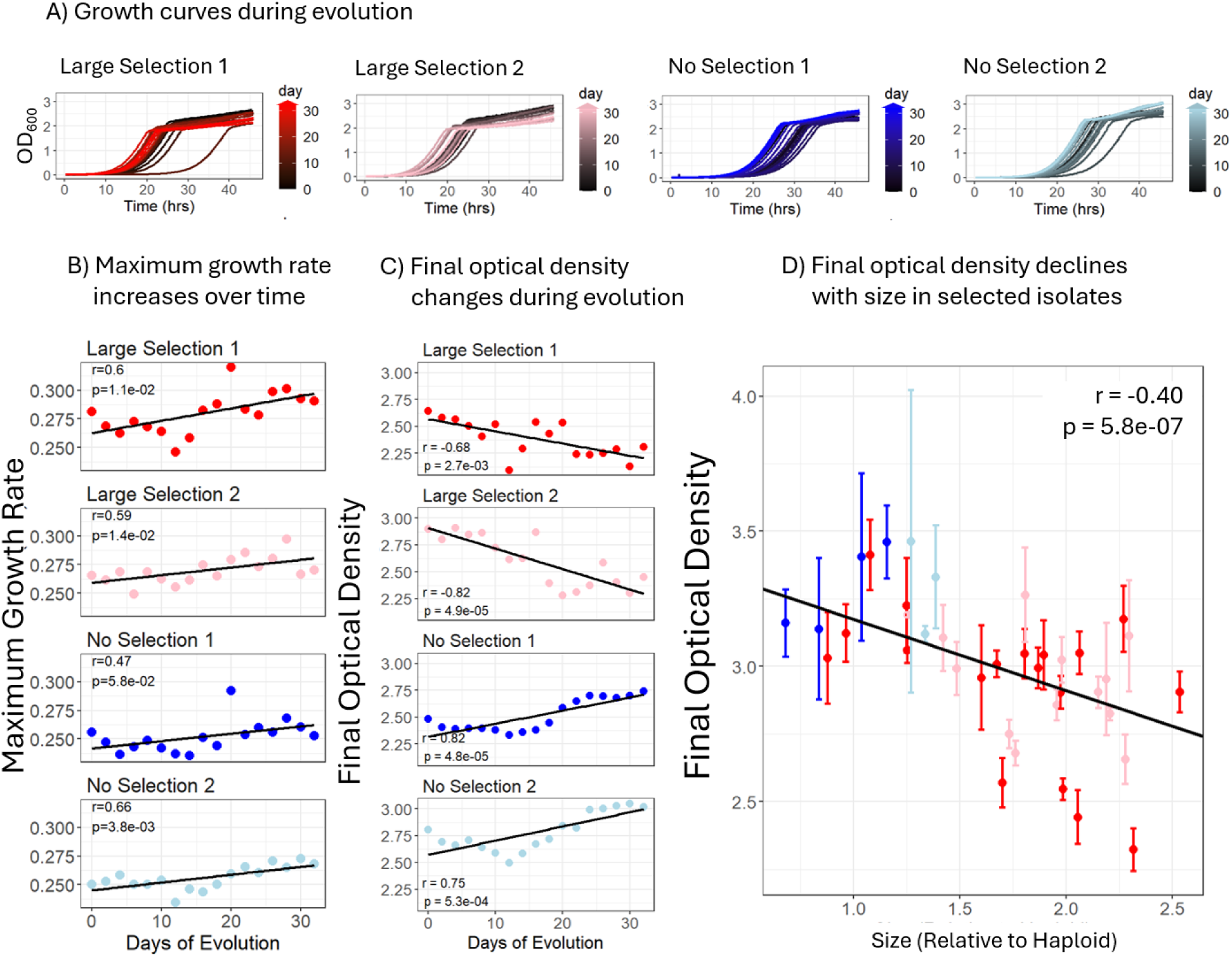
Culture optical density decreases over time in large size selection (LS) experiments but increases in the no size selection (NS) experiments. A) Plots show the raw growth curves for each 48-hour incubation period of the evolution experiment corresponding to days 0 through 32, with optical density shown on the vertical axis and hours past inoculation on the horizontal axis. Each growth curve is colored to correspond to when it was sampled during the course of the evolution, from day 0 (black) to day 32 (brightly colored). In the leftmost two plots, the black growth curves achieve higher optical density than the colored curves, consistent with panel C showing that maximum optical density decreases over the course of LS experiments. B) Maximum growth rates (hr^-1^) increase over time in all four evolution experiments. Plots show maximum log-linear growth rates plotted against days of the evolution experiment. C) Final culture (optical) density decreases over time in both LS replicates but increases for both NS replicates. The final optical density is calculated by taking the average optical density across the last 10 OD measurements of each 48-hour window, plotted against days of the evolution experiment. D) Final culture (optical) density values for randomly selected single-cell isolates decrease with the average cell size of each isolate. This plot shows the average culture (optical) density from up to 12 replicate growth curves generated for each isolate, versus that isolate’s cell size measured by forward scatter area. In panels B – D, r is the Pearson’s correlation coefficient in the regression analysis; p is the associated P-value.

To investigate different aspects of cellular physiology, we also studied the OD near the end of each growth cycle. Specifically, we estimated final culture OD as the mean of the final ten OD measurements, corresponding to the final 1.67 h of each 48-h incubation. The result revealed a clear difference between the LS and NS populations. Final OD increased over evolutionary time in both NS populations but declined progressively in both LS populations (**Fig. 2C**). The complete growth curves showed the same opposing trajectories: later NS cultures reached increasingly higher final optical densities, whereas later LS cultures reached increasingly lower final optical densities (**Fig. 2A**). Thus, cells selected for increased size continued to grow rapidly but reproducibly produced cultures with lower final OD.

Because each evolving population contained many genetically distinct barcoded lineages, we next asked whether the relationship between cell size and final OD was also apparent among clonally propagated evolved isolates. For isolates collected from both selection regimes at multiple evolutionary timepoints, we measured both growth curves and mean FSC-A. Across these isolates, larger cell size was associated with lower final culture OD (**Fig. 2D**). We then expanded the optical density analysis to an additional 86 isolates (46 from LS1 on Day 38 and 40 from its ancestor). Isolates from the ancestor population consistently reached higher final ODs than isolates from the LS population (**Fig. S1**). This larger validation set therefore confirmed that the LS selection regime experienced a decline in final OD that was retained among independently propagated clones. Together, these results show that larger cells tended to reach lower final culture densities and that the lower-OD phenotype was broadly reproducible among isolates from the LS populations.

This result showed that the relationship between cell size and cellular material had changed in the LS experiment, but OD could not reveal the exact mechanism. OD measures how strongly a culture scatters light; it does not directly measure cell number, total biomass, or demographic carrying capacity. Larger cells may scatter light differently from smaller cells, and scattering can also change with cell shape, aggregation, and internal composition. Indeed, isogenic yeast strains that differ in ploidy and apparent diameter also differ in both final OD₆₀₀ and the relationship between OD₆₀₀ and cell abundance (Fukuda 2023). The lower final OD of the LS populations could therefore indicate fewer cells, less accumulated biomass, altered scattering by individual cells, or some combination of these effects. To determine what had changed within the enlarged cells themselves, we next measured biomass across several OD values in an evolved LS population and its ancestor, revealing the opposite trend. The evolved LS population produced more biomass than the ancestor at similar OD values (**Fig S2A**). This led us to next investigate the ability of individual cells to scatter light, which is related to their internal density.

### Internal density declines as cells evolve larger size

Cellular density, the amount of intracellular material per unit volume, is normally maintained within a considerably narrower range than either cell mass or cell volume, indicating that cells generally coordinate the production of proteins and other cellular contents with expansion of cellular volume (Godin et al. 2010; Neurohr and Amon 2020). If material production kept pace as cells evolved larger size, internal density should remain relatively constant. If cellular enlargement outpaced material production, larger cells should instead become more dilute.

We measured internal density using optical diffraction tomography, allowing us to reconstruct images of living cells and measure their refractive indices (Brown et al. 2025). Refractive index describes how strongly material bends light and generally increases as the concentration of intracellular material increases. We therefore used the mean refractive index across cellular cross-sections as a proxy for internal material density.

We first analyzed samples collected at the end (day 38) of the experiment from both LS populations, both NS populations, and the unevolved ancestral population (day 0). Each evolved sample contained a genetically heterogeneous mixture of barcoded lineages and a broad distribution of cell sizes. Across all imaged cells, cellular cross-sectional area was negatively correlated with mean internal refractive index (**Fig. 3A**). In other words, larger cells tended to have lower internal densities. This result was not caused only by average differences among experimental populations: the same negative tendency was detectable among cells within each sampled population (**Fig. 3A**, insets).

**Fig 3.**
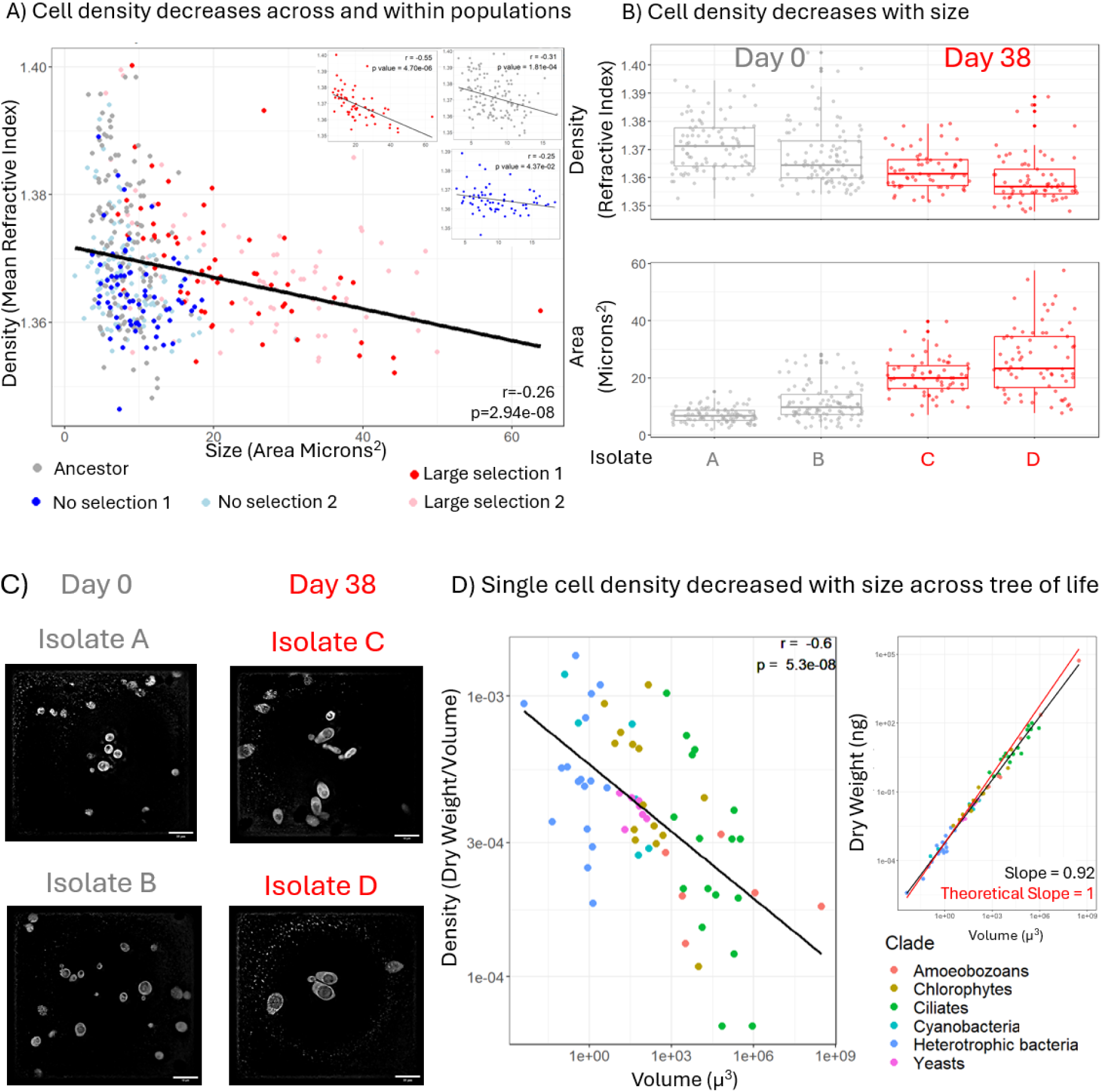
Single cell internal density decreases with size. **A)** The density of a single cell, measured *via* its internal refractive index, decreases with cell size both across all evolved populations and within each evolved population. All plots show the mean internal refractive index across all pixels corresponding to a given cell, plotted against the cell’s cross-sectional area. Colored points correspond to cells sampled from the last timepoint of the respective evolution experiment, while grey points correspond to cells sampled from the ancestral population (before evolution). No matter where we look, cell density inversely correlates with cell size. The value of r is Pearson’s correlation coefficient in the regression analysis; p is the associated P-value. **B)** When we randomly sample and propagate a single cell from the ancestral population (grey) or the evolved large population (red), we see concomitant differences in not only cell size, but also cell density. Top box plot shows distributions of each selected isolate’s internal refractive index across 4 isolates. Bottom box plot shows distributions of each selected isolate’s cross sectional area across the same 4 isolates. **C)** Micrographs of the isolates shown in **panel B**, showing both the increase in cell size and decrease in cell density from day 0 to day 38, with a 10-micron bar for reference. **D)** The decreasing cell density associated with increased cell size appears common across the tree of life. **Left**, density of 68 diverse taxa plotted against respective volume measurements, with a linear regression plotted. **Right**, Dry weight of the same 68 diverse taxa plotted against volume highlights that cell mass (dry weight) does not keep up with cell volume because the slope of the best fit line is less than 1. The black line is the line of best fit with slope labeled. The solid red line represents a theoretical 1:1 scaling relationship between dry weight and volume. Data are replotted from previous work (Lynch et al. 2022).

These single-cell measurements clarify the meaning of the final-culture OD result from **figure 2**. Larger cells likely scattered light differently because their interiors had lower average refractive indices, consistent with lower concentrations of cellular material. Thus, production of intracellular material did not fully keep pace as cellular size increased.

The shape measurements provide important context for this result. On average, the LS populations became more elongated rather than simply expanding proportionally in all dimensions. As described above, this geometry could limit the increase in internal volume associated with becoming larger, therefore reducing the amount of additional cellular material needed to maintain density. Nevertheless, larger cells still had lower internal densities. Any geometric compensation provided by elongation was therefore incomplete: cellular enlargement still outpaced the accumulation of intracellular material.

To determine whether the size–density relationship persisted after individual cells were isolated and clonally propagated, we examined four isolates selected to span differences in cell size: two derived from the ancestral population and two derived from the final timepoint of the LS population. The larger evolved isolates had greater cross-sectional areas and lower mean refractive indices than the ancestral isolates (**Fig. 3B, C**). Lower density can therefore be a stable property of clonally propagated evolved genotypes rather than only a transient feature of heterogeneous populations.

Microscopy also revealed prominent structures that resemble vacuoles in the largest isolates (**Fig. 3C**). The vacuole is an aqueous organelle with a lower refractive index than the surrounding cytoplasm, consistent with a lower concentration of cellular material (Pirone et al. 2024). Importantly, vacuole volume is known to increase disproportionately as budding yeast cells become larger. Vacuolar volume scales with cell volume with an exponent of approximately 1.3, such that larger cells devote an increasing fraction of their volume to the vacuole (Chan and Marshall 2014). Expanding the vacuole could therefore allow a cell to become larger without producing a proportional amount of new cytoplasmic material. Vacuolar expansion may consequently represent one mechanism by which cellular volume outpaces material accumulation.

We next asked whether the negative relationship between size and density was specific to our evolved yeast or reflected a broader biological pattern. Previous studies have largely examined density homeostasis within a species, as cells grow or are experimentally perturbed (Bryan et al. 2010; Godin et al. 2010; Odermatt et al. 2021; Liu et al. 2022). However, direct measurements across 12 bacterial and archaeal species have also revealed sublinear scaling of dry mass with cell volume, such that larger species have lower dry-mass density (Khachikyan et al. 2019). To ask whether this relationship extends more broadly across the tree of life, we analyzed dry-mass and volume measurements compiled for 68 species spanning major phylogenetic groups (Lynch et al. 2022). Across these taxa, cellular density, calculated as dry mass divided by volume, was negatively correlated with cell volume (**Fig. 3D**). Dry mass still increased as cells became larger, meaning that large cells contained more total cellular material. However, mass increased more slowly than volume, so larger cells contained less material per unit volume.

Together, the growth, morphology, single-cell imaging, isolate, and comparative analyses distinguish density homeostasis during growth from density scaling over evolution. Cells selected for increased size preserved basic growth performance: their maximum growth rates did not decline and instead increased during evolution. However, they did not preserve the ancestral relationship between cell size and material concentration. Even after cells reproducibly altered their average geometry, intracellular material failed to increase in proportion to cellular enlargement. Whereas evolutionary miniaturization can establish a stable new size with limited fitness costs (Garoña et al. 2026), evolution toward larger size in our experiment produced a stable shift toward lower internal density. The cross-species analysis extends this result, showing that density can be maintained within cell types while declining systematically with cell size across species.

### Increased cell size evolves through multiple genetic routes

The inverse relationship between cell size and density would be less general if every LS cell descended from one unusual mutant that increased size through a single mechanism. A central advantage of the barcoded evolution platform is that it began with many independently evolving lineages, many of which persist through the end of the evolution experiment (**Fig S3**). We surveyed these diverse lineages to ask whether selection for increased size recruited different genetic solutions across adaptive isolates or was dominated by similar patterns of genetic change.

We first examined DNA content because ploidy and cell volume scale approximately linearly in budding yeast (Turner et al. 2012). Increased ploidy, the number of complete genome copies within a cell, is a well-established route to increased yeast cell size (Chan and Marshall 2010). We stained fixed isolates collected on days 0, 18, and 38 with SYTOX Green and compared their DNA fluorescence and FSC-A values with those of haploid and diploid controls. DNA content was positively associated with cell size across the evolved isolates (**Fig. 4A**). At the beginning of the experiment, isolates from all populations resembled the haploid control in both size and DNA content. By day 18, many LS isolates had DNA content and FSC-A values similar to those of diploid controls, whereas NS isolates remained predominantly haploid-like. By day 38, LS isolates displayed broad variation in both size and DNA content. This suggests that increased ploidy contributed substantially to the response to size selection but did not account for all of the observed size variation.

**Fig 4:**
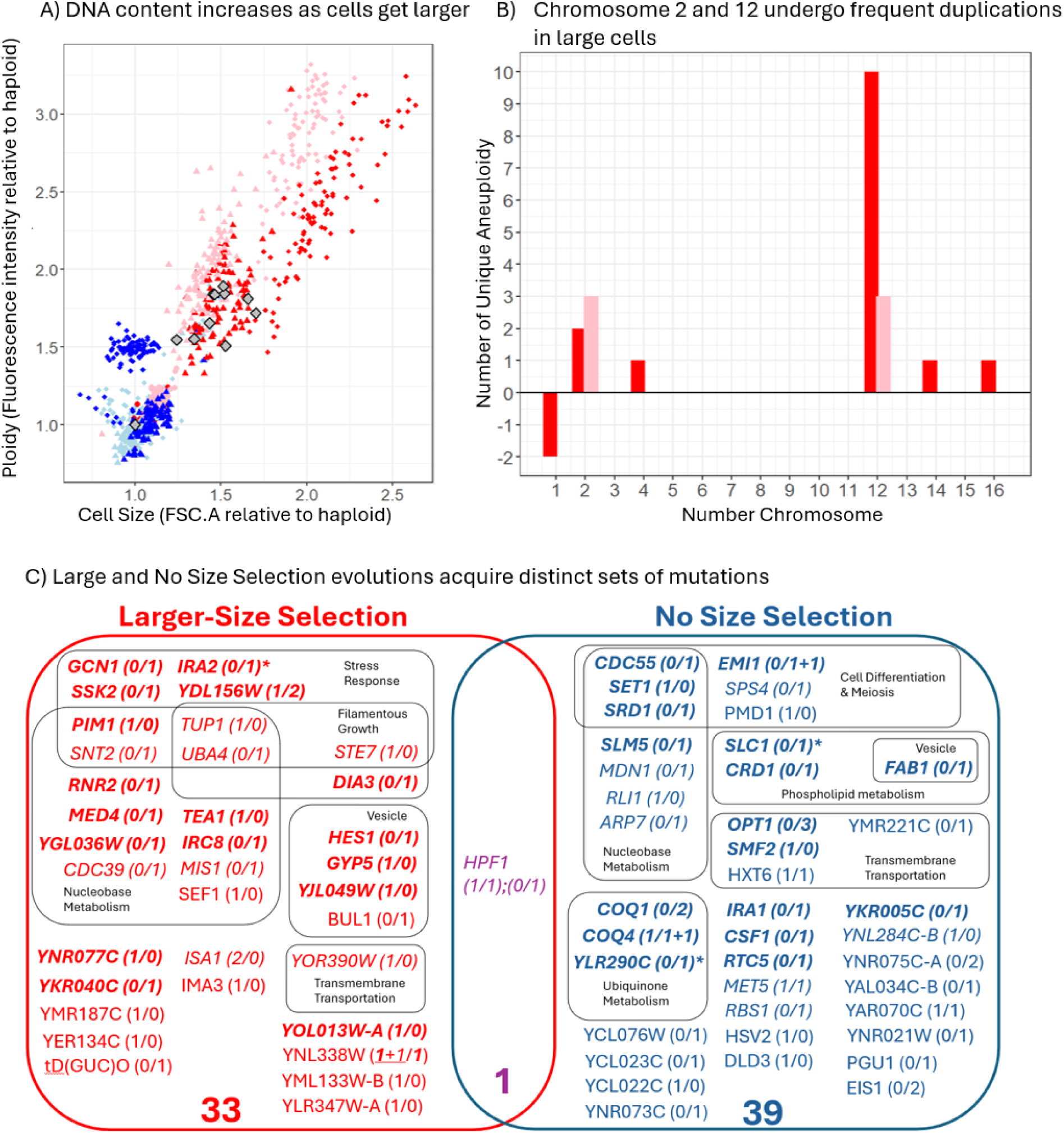
There are diverse ways to increase cell size. **A)** The DNA content of isolates, measured via SYTOX Green fluorescence emission, correlates with isolate cell size, measured via forward scatter area, across evolved isolates. All measured values are reported relative to a haploid control. At day 0 of all experiments, cells have similar size and DNA content to a haploid control (red and blue circles cluster near the gray diamond at coordinates 1,1). While cells from the NS experiments (blues) remain like the haploid control, cells from the LS experiments (reds) increase in size and ploidy. For example, by day 18, the cells under selection for large size cluster near the diploid controls (the red/pink triangles cluster near the other grey diamonds). By day 38, cells from the LS experiments are larger and have greater DNA content than diploids (the red/pink diamonds are spread on the top right of the graph). **B)** Whole genome sequencing reveals diverse aneuploidy present across cell lineages that underwent large size selection. Among these lineages, chromosome 12 and chromosome 2 experience the most duplication across sequenced isolates. **C)** Genes associated with single-nucleotide mutations detected in isolates with unique barcodes at the end of size selection experiments. In total, 33 genes accumulated mutations in at least one isolate by the end of the large-size selection experiments; 39 genes did so during the no size-selection experiments. The gene ontology (GO) terms for mutated genes are noted with colored circles and simplified description. Except for HPF1, there is little overlap in the mutated genes across selection pressures, though there is more overlap within each pressure. For example, under large size selection, YDL156W was found to possess a mutation in one isolate from replicate one and two isolates from replicate 2. (YDL156W is involved in the DNA damage response). The numbers in the parenthesis indicate the number of unique barcodes found possessing a mutation in a given gene, separated by replicate. The genes with italic texts indicate synonymous mutations. The genes with italic and bold texts indicate nonsynonymous mutations. The genes with italic texts and an asterisk (*) indicate nonsense mutations. The genes with regular texts indicate mutations within the upstream or downstream 5kb of the genic regions or non-coding genes. When more than one type of mutation is associated with the same gene, we use the plus symbol (+) to indicate it. For example, under large size selection, “YNL338W (***1***+*1*/***1***)” indicates that the gene YNL338W was found mutated in a total of 3 isolates: one in replicate one had a nonsynonymous mutation, a second in replicate one had a synonymous mutation, and a third in replicate two had a nonsynonymous mutation. YNL338W is a putative uncharacterized protein.

Additional genome copies could help larger cells preserve composition because the ploidy-to-cell-size ratio is a key determinant of proteome scaling; disproportionately large yeast cells exhibit genome dilution and a starvation-like proteome (Lanz et al. 2024). More generally, increasing ploidy can extend the range of cell sizes over which biosynthesis scales with cell volume by increasing the number of gene copies available for transcription (Cadart and Heald 2022). At the other extreme, experimentally induced endopolyploid yeast reached 16C–32C DNA content, but ploidy-associated enlargement created cell-surface stress that limited viability (Barker et al. 2025). Increased ploidy may therefore expand biosynthetic capacity without guaranteeing proportional scaling of all cellular material.

Moving beyond whole genome duplication, we next used whole-genome sequencing of 87 isolates to identify changes in the copy number of individual chromosomes. Aneuploidies, gains or losses of particular chromosomes rather than the entire genome, were common among the 70 sequenced LS isolates. Chromosomes II and XII showed the most frequent duplication events, with five and twelve gains, respectively (**Fig. 4B**). No aneuploidy was detected among the 17 sequenced NS isolates. The repeated gain of chromosome XII is notable because this chromosome contains the ribosomal DNA array and genes involved in vacuolar and membrane-associated processes. These functions could plausibly affect the ability of cells to produce material or reorganize their internal architecture as they enlarge.

Sequence-level mutations also differed markedly between the selection regimes. Mutations associated with 33 genes were detected only among LS isolates, whereas mutations associated with 39 genes were detected only among NS isolates; *HPF1* was the only gene mutated in both conditions (**Fig. 4C**). Interestingly, genes mutated under large-size selection included functions related to vesicle-mediated transport, stress response, and filamentous growth; these functions are rarely or not seen in the genes mutated in the NS experiments. These functions are relevant to the movement of material within cells and to the elongated morphology observed in the LS populations (Pruyne et al. 2004; Cullen and Sprague 2012).

Selection for increased size therefore recruited several kinds of genetic change, including increases in whole-genome DNA content, gains of individual chromosomes, and sequence-level mutations affecting diverse cellular functions. This response differs from recently evolved miniature yeast, in which substantial decreases in size arose through changes in cell-cycle and nutrient-sensing pathways without corresponding changes in ploidy (Garoña et al. 2026). Together, the studies show that yeast cell size can evolve in either direction through distinct regulatory and genomic routes.

The genetic results demonstrate that the population-level relationship between increased size and lower density emerged in genetically heterogeneous populations rather than in a population defined by one engineered large-cell mutation. Multiple evolving lineages found different ways to become larger, yet the populations as a whole repeatedly displayed the same central architectural tendency: cellular enlargement exceeded the maintenance of intracellular material concentration.

### The evolved large-cell state is associated with stable, environment-dependent tradeoffs

The preceding results identified a reproducible departure from density homeostasis. The LS populations maintained rapid growth but evolved larger cells whose intracellular material did not increase in proportion to cell size. If maintaining intracellular material concentration is important for robust cell function, the consequences of this change may become more apparent when cells are challenged outside the environment in which they evolved. We therefore asked whether the LS and NS populations responded differently to two environmental perturbations that challenge cellular organization in different ways: elevated temperature, which increases demands on proteostasis and reorganizes cellular architecture, and hyperosmotic salt, which removes water, compresses cellular volume, and increases intracellular molecular crowding (Miermont et al. 2013; Babazadeh et al. 2013; Wallace et al. 2015; Keuenhof et al. 2022).

We revived frozen samples from both evolved LS populations and both evolved NS populations, regrew them through saturation, and standardized each population to the same starting cell number before measuring three replicate growth curves per evolved population under each condition. We compared growth at 27°C, 30°C, and 35°C and at NaCl concentrations of 0, 0.2, and 0.4 M. OD₆₀₀ was measured every ten minutes throughout each growth curve. Replicate cultures produced similar growth curves and consistently showed the same condition-dependent shifts, allowing us to distinguish changes in the LS population from changes in the NS population (**Fig. 5 & Fig S4**).

**Figure 5.**
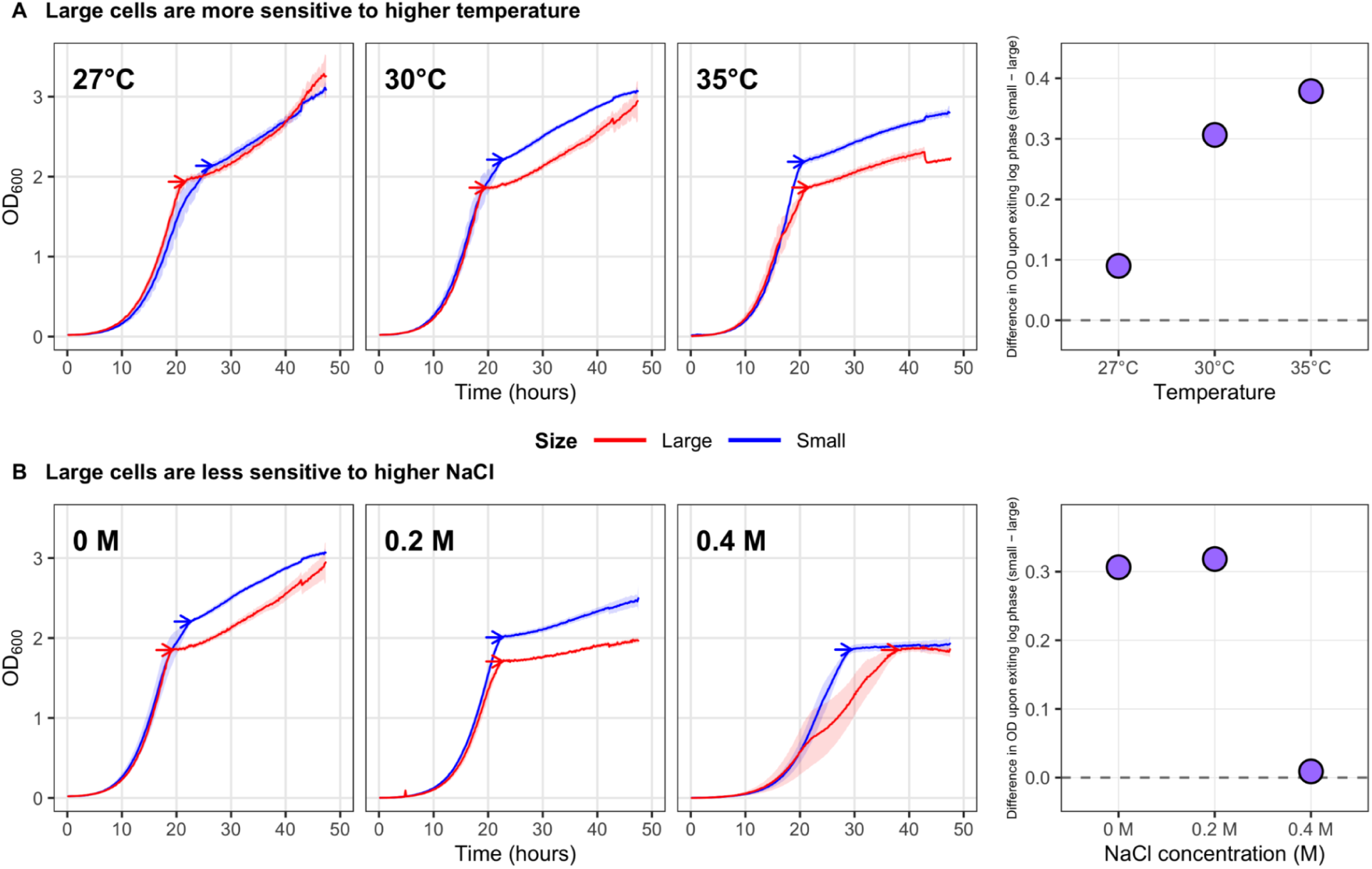
Evolved large-cell populations show distinct OD responses to temperature and salt. Frozen samples representing the final timepoint (Day 38) of the four evolution experiments were revived and grown to saturation. Each independently evolved population was then divided into three replicate cultures, for a total of six corresponding to LS and six to NS, from which growth curves were collected in various environments by measuring OD_600_ every ten minutes. Lines show the mean OD across the six replicate cultures with standard error across replicates shown as a ribbon. NS populations are shown in blue and LS populations in red. Arrows on the growth curves indicate the transition from exponential to post-exponential growth, at which OD was quantified. Each purple point shows the magnitude of the mean difference between the LS and NS populations at this transition; higher purple points therefore indicate greater separation between the selection regimes. **(A)** At 27°C, LS and NS populations reached approximately the same OD at the end of exponential growth. The NS populations changed comparatively little as temperature increased, whereas the LS populations reached progressively lower OD₆₀₀ values.

The response to temperature was driven mainly by a change in the LS population. At 27°C, the LS and NS populations reached approximately the same OD at the end of exponential growth. The NS populations reached a similar OD as temperature increased, whereas the LS populations reached progressively lower OD values at 30°C and 35°C (**Fig. 5A**). The difference between the NS and LS populations therefore widened with increasing temperature because the LS populations were more sensitive to elevated temperature.

Both populations responded negatively to increasing NaCl, but the decline was more pronounced in the NS populations. In the absence of added NaCl, the NS populations reached an OD approximately 0.3 units higher than the LS populations (**Fig. 5B**). As NaCl increased, this difference narrowed because NS OD declined more sharply than LS OD. At 0.4 M NaCl, both populations reached approximately the same OD after 48 hours of growth. Salt therefore did not improve growth of the LS populations relative to its own unstressed condition. Instead, the LS populations were less strongly affected by salt than the NS populations by this culture-level measure. Because osmotic compression increases intracellular molecular concentration and can slow diffusion and cellular processes in crowded yeast cells (Miermont et al. 2013; Babazadeh et al. 2013), the reduced sensitivity of the LS populations to salt is consistent with the possibility that their lower starting density provides greater tolerance for osmotic compression.

Together, these assays reveal stable but opposing environmental responses associated with the evolved large-cell state. These results show that the physiological consequences of selection for increased size depend on environmental context. They are consistent with the possibility that failure to maintain proportional intracellular material concentration creates conditional tradeoffs. The most direct conclusion is that the evolved LS state, which includes changes in size, density, geometry and ploidy, was associated with greater sensitivity to elevated temperature but lower sensitivity to hyperosmotic salt.

The increasing separation between NS and LS at higher temperatures was therefore driven primarily by a decline in the LS populations. **(B)** OD at the end of exponential growth decreased with increasing NaCl in both NS and LS populations. This decline was more pronounced in the NS populations, causing the approximately 0.3-unit difference observed without added NaCl to narrow as salt increased and to disappear at 0.4 M NaCl, where both population types reached similarly low OD values.

## Discussion

Selection for increased cell size repeatedly produced cells in which volume expansion outpaced the accumulation of cellular material. This outcome was striking because density itself was never directly selected: selection acted on apparent cell size, and larger cells arose through diverse genetic and morphological routes, including altered geometry, changes in ploidy, aneuploidy, and sequence-level mutations. Nevertheless, lower internal density emerged repeatedly and persisted in clonally propagated isolates. The same qualitative relationship was also evident across species, where cellular mass increased with volume but less than proportionally. Thus, reduced density may be more than an incidental defect of a particular oversized mutant. It may be a predictable correlated consequence of evolutionary increases in cell size.

This possibility changes how density homeostasis can be viewed. Within proliferating cells, mass and volume are remarkably well coordinated: density varies little through ordinary growth and is resistant to many perturbations that alter biosynthesis (Bryan et al. 2010; Odermatt et al. 2021; Liu et al. 2022). Our results suggest that homeostasis need not imply an evolutionarily fixed density set point. Instead, cells may robustly maintain one relationship between mass and volume over short timescales while evolution gradually establishes another. An important unanswered question is therefore whether the evolved LS cells actively defend their lower density. For example, following acute osmotic or biosynthetic perturbations, do they return to a density characteristic of the evolved genotype rather than that of the ancestor? If so, density would be better understood as an evolvable physiological state than as a fixed cellular constant.

Evolution toward larger versus smaller cell size may also face fundamentally different scaling constraints. Previous work (Garoña et al. 2026) evolved dramatically miniaturized yeast that retained robust growth, biomass production, and ribosome biogenesis without increasing genome copy number, largely through coordinated changes in nutrient sensing and cell-cycle regulation. At the opposite extreme, very small prokaryotes can contain disproportionately large amounts of material per unit volume, possibly because a minimum complement of DNA, enzymes, membranes, and other machinery cannot decrease indefinitely with size (Khachikyan et al. 2019). Enlargement presents the converse problem: each additional unit of volume creates new demand for biosynthesis. Consistent with this asymmetry, increased DNA content was common among our large evolved isolates. Because additional genome copies can extend the size range over which transcription and biosynthesis scale with volume (Neurohr et al. 2019; Cadart and Heald 2022), increased ploidy during large-size evolution may serve not only as a route to becoming larger, but also as partial compensation for the biosynthetic demands created by enlargement. The persistence of reduced density despite increased DNA content suggests that this compensation was incomplete. Testing whether polyploidization limits the magnitude of dilution, rather than simply causing enlargement, could help distinguish these possibilities.

Reduced density need not have uniformly negative physiological consequences. Both ends of the concentration spectrum can interfere with cellular function. Excessive enlargement and cytoplasmic dilution impair gene expression, signaling, and cell-cycle progression, whereas osmotic compression and increased molecular crowding reduce protein mobility and slow numerous cellular processes (Miermont et al. 2013; Neurohr et al. 2019; Neurohr and Amon 2020). Density may therefore define a biophysical axis along which different environments favor different cellular states. Our stress experiments are consistent with this possibility. The large evolved populations were more sensitive to elevated temperature, yet their growth was less strongly affected by increasing NaCl than that of the NS populations. Because hyperosmotic conditions reduce cellular volume and increase intracellular concentration, one possibility is that a more dilute starting state provides greater capacity to tolerate osmotic compression before excessive crowding is reached. We did not directly measure intracellular density under salt stress, so this mechanism remains speculative. The greater temperature sensitivity of the LS populations is also consistent with the widespread temperature–size relationship in unicellular organisms, in which cells tend to be smaller at higher growth temperatures (Forster et al. 2013; Zakhartsev and Reuss 2018). More generally, these opposing responses to elevated temperature and salinity suggest that evolution of cell size and density can reshape environmental tradeoffs rather than simply impose a universal fitness cost.

What physically creates the additional low-density volume remains an important mechanistic question. The conspicuous vacuole-like structures in large evolved cells provide one possibility, particularly because budding-yeast vacuoles superscale with cell volume (Chan and Marshall 2014). Increasing the fraction of the cell occupied by a relatively dilute compartment could allow substantial enlargement without proportional production of cytoplasmic macromolecules. Other contributions could include changes in water, ions, osmolytes, organelle composition, or the concentrations of particular classes of macromolecules. Our refractive-index measurements report an average cellular property and therefore cannot yet distinguish dilution of the cytoplasm itself from redistribution of volume into low-density compartments. Likewise, the cross-species relationship cannot by itself determine whether lower density in larger taxa reflects a shared physical constraint, repeated adaptation, or correlated ecological and phylogenetic differences.

Experimental evolution supplies a complementary piece of evidence by showing that the relationship can emerge rapidly within a single species. If the same principle proves general, the decrease in density observed across the tree of life may not simply represent failure of cells to maintain an optimal state. Instead, accepting a more dilute internal architecture, and compensating through changes in genome content, organelle organization, or physiology, may be one of the ways evolution makes larger cells possible.

## Materials and Methods

### Base strain

All *S. cerevisiae* lineages originated from a barcoded pool derived from strain SHA185, described previously (Levy et al. 2015), with the following genetic background: MATɑ, ura3Δ0, ybr209w::Gal-Cre-KanMX-1/2URA3-loxP-Barcode-1/2URA3-HygMX-lox66/71.

### Base media

All size selection experiments were conducted in ‘M3’ media, a glucose-limited media lacking uracil as defined previously (Levy et al. 2015; Schmidlin et al. 2024).

### Initiating the size selection evolution experiments

All evolution experiments started with the same pool of approximately 300,000 uniquely barcoded yeast lineages as defined in Schmidlin et al. 2024 (Schmidlin et al. 2024). To begin the experiment, a pea sized amount of frozen yeast barcode library was grown in 6 mL YPD for 4 h at 30℃ in a shaking incubator at 220 rpm. One million cells were transferred from this culture to each of four tubes that served to found four evolution experiments.

### Performing the size selection evolution experiments

Four independently evolving populations were propagated in parallel: two Large Size Selection (LS) populations and two No Size Selection (NS) control populations. During each transfer cycle, populations were grown for 48 h in 6 mL of M3 medium at 30°C and 70 rpm in a compact rocking incubator (TVS062CA; Advantec MFS). Culture optical density at 600 nm was recorded automatically every 10 min. After 48 h, 1 mL of each culture was vortexed vigorously for 1 min, diluted in PBS, and analyzed on a BD FACSAria flow cytometer at the Arizona State University Flow Cytometry Facility. For each LS population, one million cells from approximately the largest 10% of the forward-scatter distribution were sorted into 6 mL of fresh M3 medium. For each NS population, one million cells were sorted without regard to forward scatter and transferred into fresh medium. The populations were then grown for another 48 h, and this growth-and-sorting cycle was repeated for 20 transfers, corresponding to approximately 160 generations based on an estimated eight generations per cycle (Levy et al. 2015). After each transfer, the remaining culture was collected by centrifugation for 3 min at 5,000 rpm, resuspended in approximately 30% glycerol, divided into freezer stocks, and stored at −80°C. These archived populations were later used to quantify barcode frequencies and recover evolved isolates.

### Size Selection Evolution flow cytometry gating parameters

Flow cytometry gating was performed using 3 parameters, Forward Scatter Area (FSC.A), Forward Scatter Height (FSC.H), and Side Scatter (SSC.A). For each condition independently, a sample was taken to plot parameter SSC.A against parameter FSC.A. Gating was used to filter events that clustered around 0,0. Within this gate, parameter FSC.H was plotted against parameter FSC.A and a second gate was performed to select for single cells. Using the second gate, a distribution based on the FSC.A parameter was used to sort the 1 million transferred cells. For the Large Size Selection experiments, the first 1 million events to be recorded within the ∼top 10 % of the second gate’s FSC.A distribution were sorted and propagated. For the No Size Selection experiments, the first 1 million events to be recorded anywhere within its sample FSC.A distribution were sorted and propagated.

### Staining clonally propagated isolates with SYTOX Green

We measured DNA content in clonally propagated isolates using SYTOX Green staining, following Schmidlin et al. (2024) (Schmidlin et al. 2024). Frozen population samples from selected timepoints of the Large Size Selection and No Size Selection experiments were first streaked onto YPD agar and incubated at 30°C for 48 h. Each colony that formed on the plate was assumed to have arisen from a single cell and therefore represented a clonally derived isolate from the evolving population. For each population sample, 94 individual colonies were picked with sterile toothpicks and transferred separately into wells of a 96-well plate containing 150 µL M3 medium. The remaining two wells contained haploid and diploid control strains grown in 150 µL YPD. Plates were incubated at 30°C for 48 h, allowing each colony-derived isolate to expand into a clonal culture.

After growth, 75 µL from each clonal culture was transferred to a second 96-well plate containing 75 µL of 30% glycerol and stored at −80°C. The remaining cells in the original plate were prepared for DNA-content measurement. Plates were centrifuged at 4,500 rpm for 1.5 min, the supernatant was removed, and cells were fixed in 150 µL of 70% molecular-grade ethanol. The plates were centrifuged again, and the ethanol was removed. Fixed cells were treated with 50 µL RNase A at 2 mg/mL for 2 h at 37°C to remove RNA. Samples were then centrifuged, the supernatant was discarded, and 20 µL pepsin at 5 mg/mL was added to each well. Cells were incubated with pepsin at 37°C for 20–45 min, centrifuged, and cleared of supernatant. Finally, cells were resuspended in 149 µL Tris-HCl buffer (50 mM, pH 8.0) containing 1 µL of 1 mM SYTOX Green.

### Flow-cytometric measurement of DNA content and cell size

DNA content and cell size were measured using a Thermo Fisher Attune NxT flow cytometer at the Arizona State University Flow Cytometry Core Facility. Each sample consisted of many cells descended from one colony and therefore represented a clonally propagated isolate rather than a heterogeneous evolving population. Events were first gated using forward-scatter area and side-scatter area to exclude debris and events near the origin. A second gate based on forward-scatter area and forward-scatter height was then used to retain single-cell events and exclude aggregates. For the gated singlet cells from each clonal isolate, forward-scatter area was used as a proxy for cell size, and SYTOX Green fluorescence in the BL1-A channel was used as a proxy for DNA content. The mean forward-scatter and fluorescence values across the measured cells were used to estimate the characteristic cell size and DNA content of each isolate.

### Measuring growth curves of clonally propagated isolates

To measure growth phenotypes of individual evolved genotypes, frozen population samples from days 8, 18, and 38 of the Large Size Selection and No Size Selection experiments were first streaked onto YPD agar. After colonies formed, individual colonies were picked with sterile toothpicks and transferred separately into liquid medium. Because each colony was assumed to have arisen from a single cell, each colony-derived culture was treated as a clonally propagated isolate from the corresponding evolving population. Isolates were selected at random from the available colonies to capture variation in cell size across selection conditions and evolutionary timepoints. Frozen stocks of these clonal isolates were revived in 6 mL M3 medium and incubated at 30°C for 24 h. Cell abundance was then measured, and approximately 1.5 × 10⁶ cells from each revived isolate were transferred into 6 mL of fresh M3 medium to initiate a growth-curve assay. Cultures were incubated for 48 h at 30°C and 70 rpm in a compact rocking incubator (TVS062CA; Advantec MFS). Optical density at 600 nm was recorded automatically every 10 min. Each isolate was assayed in 2–12 independent replicate cultures.

### Measuring Maximum Growth Rate per Growth Cycle

In order to measure the maximum growth rate per growth cycle, specific growth rates were calculated by taking the log linear slope of a sliding window of 18 timepoints (3 hours) across each growth curve. Following blanking, to avoid measurement noise recorded at low OD_600_ readings, all values were shifted up by the difference between 0.001 and the lowest growth curve OD_600_. Linear regressions with an R^2^ greater than 0.99 were kept, with the largest slope being identified as the maximum growth rate for each respective growth curve.

### Calculating Final OD_600_ Near the End of Each Growth Cycle

In order to reduce noise in OD measurements during the stationary phase, the final OD per growth cycle was measured by taking the average of the last 10 OD_600_ readings (∼1.67 hours) of a growth curve.

### Measuring Refractive Index Using Optical Diffraction Tomography

We grew each sample to saturation in M3 at 30℃ for 48 hours and then performed a 1:100 dilution. We pipetted 40ul of each solution onto a 40mm diameter round no 1.5 cover glass, and dropped a square 25 mm cover glass on top. We then placed a few drops of water on top of the 25mm cover glass to support imaging with a water immersion objective. We imaged the samples using Fourier synthesis optical diffraction tomography (FS-ODT) in non-multiplexed mode with 785nm illumination on an instrument using a 100x NA 1.3 oil immersion objective (Olympus UPlanFL N) to project plane waves and a 60x NA 1.0 water immersion objective (Olympus LUMPLFLN60XW) to image the collected light onto a camera with 18um pixels (Phantom VEO-1010L-72G-M). Further details of the instrument are described in previous work (Brown et al. 2025). At each position in the sample, we collected tomographic data using 349 angles with an exposure time of 0.5ms per angle. We ensured that each data set included a cell-free background image, which is required for the reconstruction algorithm. We reconstructed the 3D refractive index on a 21×960×1040 grid with voxel size 1um × 0.075 × 0.75um using custom python code implementing the beam propagation method (BPM) as a forward model, and the fast iterative shrinkage-thresholding algorithm (FISTA) for the inverse solver (Beck and Teboulle 2009). We imposed constraints that the real part of the refractive index must be larger than the background index of 1.333 and that the imaginary part of the refractive index is zero. For regularization, we adopted the plug-and-play prior approach using a 1×3×3 median filter as a proximal operator (Venkatakrishnan et al. 2013).

### Analyzing Samples for Refractive index and Size

Microscopy images for measuring refractive index were analyzed using Fiji software on ImageJ. To measure the refractive index for cells within each microscopy image, we picked a single axial plane where the Z layer showed the largest cross-sectional area across all cells. Cells were selected by hand using the polygon tool to encompass their area, excluding cells which showed poor resolution. Each cell’s encompassed area was recorded for its size measurement and the average refractive index for the encompassed area was used as a proxy measurement for the cell’s internal density.

### Measuring Dry Mass of Populations

Dry mass was measured across the growth curves of LS Replicate 1 at Day 0 and Day 38 for OD600 values of approximately 0.5, 0.75, 1.00, 1.5, 1.75, 2, in triplicate. Approximately 1.5 million cells in 6 mL of M3 media were used to start each liquid culture. Liquid cultures were incubated at 30℃ with an rpm of 70 until desired OD600 measurements were reached. When OD600 measurements were reached, the sample was removed to measure cell count and dry mass. 5 mL of each sample was transferred into their own 15 mL falcon tube, and spun down at 4,400 rpm for 1.5 minutes. The supernatant was then removed and cells were resuspended in 1 mL of a PBS buffer solution. The resuspended media was transferred to a preweighed 1.8 mL eppendorf tube and spun down at 20,000 rpm for 30 seconds. The supernatant was removed and the eppendorf tubes were placed in a heat block at 100℃ overnight. Following heating, the eppendorf tube mass was measured and single cell mass was calculated by dividing the difference in eppendorf tube mass by the sample’s estimated cell count for 5 mL of culture. We trust this approach as values from the ancestor population did not differ significantly when compared to previous work (Łabędź et al. 2017) (see **Figure S2D** which reported a single cell dry weight measurement of 47.65 +−1.05 pg across a total range of 40 to 67 pg). To measure statistical differences to previous work (Łabędź et al. 2017), we randomly generated 1000 values, using set.seed (123), to fit the measurement of 47.65 +−1.05 pg across a total range of 40 to 67 pg, and performed a student T test with Bonferroni correction.

### Microscopic imaging of the different yeast populations and image analysis

Microscopy was performed on Large and No Size Selection populations on days 0, 12, 18, 26, and 32 of the evolution experiment. Ten µL of yeast cell culture from each yeast population were set aside right before the cell sorting, fixed with 4% paraformaldehyde at room temperature for 1 hour, and washed three times with PBS. The nuclei were stained with 1 µM Hoechst (Invitrogen, USA) and cell membranes were stained with 1 μg/mL wheat germ agglutinin (WGA) conjugated with Alexa Fluor 488 (Invitrogen, USA) for 30 minutes at room temperature. The PBS-washed cells were resuspended in vectashield (Vector Labs, USA) and mounted on microscope slides.

The imaging was carried out with a 60X objective on the fluorescence Olympus VS200 Slide Scanner (Olympus, VS200, Tokyo, Japan) at the ASU microscopy core. A 5 mm x 5 mm imaging window was set to monitor the yeast population with a 5.60 µm-thick Z stacks in 0.57 µm Z spacing to obtain an extended focus imaging (EFI) projection.

Images for each population were analyzed using VS200 Desktop (version 3.4.1). Each cell was identified using the WGA-488 signal to set the threshold for the cellular boundary and the Hoechst-stained nuclear signal was used to identify single cells. Defined objects were later analyzed for the area, perimeter, mean radius, mean gray values, shape factor and sphericity, and filtered to eliminate cell aggregates, cells at the edges, irregular shapes and staining yielding from 18,000 to 78,000 cells per image.

### Measuring Density of Species across the tree of life

Data to calculate the density of different lineages across the tree of life was collected from reference (Lynch et al. 2022) Supplementary Table 1. Density calculations were performed by dividing a species average mass by its average volume.

### Identifying adaptive mutations using whole genome sequencing

To better understand genetic mutations associated with an increase in cell size, whole genome sequencing was performed on random isolates that underwent flow cytometry forward scatter analysis. Isolates were selected to capture a range of size across different timepoints (Day 0, Day 18, and Day 38 of the evolution experiment) to investigate if genetic perturbations that initially conferred an increase in cell size persisted throughout the evolution experiment.

To perform whole genome sequencing, isolates were streaked out onto YPD plates and single colonies were grown in YPD to saturation. Isolates were then treated with 250 uL of 0.1 M Na2EDTA, 1 M sorbitol, and 5 U/uL zymolase for a minimum of 15 minutes at 37C in order to remove the cell wall. Cells were then lysed by adding 250 uL of 1% SDS, 0.2 N NaOH and then inverted several times to mix the solution. Proteins and cell debris were removed with 5 M KOAc by spinning for 5 min at 15,000 rpm. The supernatant was transferred to a new tube, and DNA was precipitated with 600 uL isopropanol by spinning for 5 minutes at 15,000 rpm. The supernatant was removed and the remaining pellet was washed with 70% molecular grade ethanol, before being resuspended in 50 uL water containing 10 ug/mL RNAse. The extracted DNA was quantified using the NanoDrop spectrophotometer and samples were diluted to a concentration of 50 ng/uL for sequencing library preparation.

Sequencing libraries were made using Illumina DNA Prep kit by diluting reactions by ⅕. Samples were prepared in 6 uL volumes between 20 and 100 ng of DNA. 2 uL of both BLT and TB1 were added to the DNA and incubated on a thermocycler at 55C (lid 100C) for 15 min. Following incubation, the beads were washed twice using 20 uL of TWB. After the second wash, 4 uL of EPM, 4 uL of water, and 2 uL of UD indexes were added into each sample. Depending on the initial DNA concentration, PCR was performed following the Illumina guidelines: lid 100C, 68C for 3 min, 98C for 3 min, [98C for 45 s, 62C for 30 s, 68C for 2 min] for 6 to 10 cycles, 68C for 1 min, 10C hold. Following the aforementioned PCR steps, products were cleaned using a double side sized selection: 4 uL of each sample was pooled together (totaling 32 uL across 8 samples), and 28 uL of water plus 32 uL of SPB was added to the pooled sample. After 5 min of incubation, 25 uL of supernatant was transferred to a new tube containing 3 uL of SPB. Following the transfer, beads were washed with fresh 80% molecular grade ethanol and libraries were eluted in 12 uL RSB. Following the elution, samples were multiplexed using Illumina’s unique dual (UD) index plates (A-D) and sequencing was performed with 2×150 paired end sequencing on HiSeq X at Psomagen (Rockville MD).

### Variant Calling

Variant calling was performed using GATK as described here: https://github.com/gencorefacility/variant-calling-pipeline-gatk4. Identified variants were then annotated using SnpEff (Cingolani et al. 2012). Variant call files from 87 isolates were analyzed in R and compared to reference strain GCF_000146045.2 (Genome assembly 64: sacCer3). During analysis SNPs were ignored if all of the evolved isolates possessed them. Additionally, we also ignored SNPs that were present in a substantial number of evolved lineages, as they likely represent background mutations that were present in a substantial portion of the cells representing the landing pad strain (Levy et al. 2015).

### Identifying aneuploidy across isolates

Sequenced isolate data was used to determine if aneuploidy was present across specific chromosomes. To accomplish this the mitochondrial chromosome was ignored, and we measured the genome-wide average read depth for an isolate. As aneuploidy could skew the genome wide average read depth, the three chromosomes which deviated the most from the average were ignored for a second calculation. Using this second average a Student’s t-test was used to measure if the average read depth for each chromosome, excluding the mitochondrial chromosome, fell outside a 95% confidence interval. Chromosomes with an average read depth falling outside the 95% confidence interval were deemed as aneuploid.

### Gene Ontology Analysis

The gene ontology (GO) terms for the gene are annotated by gProfiler. Only Biological Process terms were used. The choice of representative terms was based on the groups with statistics showing higher enrichment and higher number of genes per term. The representative terms plotted in Figure 4 include GO:0006950 (response to stress), GO: 0036267 (invasive filamentous growth), GO:0016192 (vesicle-mediated transport), GO:0055085 (transmembrane transport), GO:0030154 (cell differentiation), GO:0051321 (meiotic cell cycle), GO:0006139 (nucleobase-containing compound metabolic process), GO:0006644 (phospholipid metabolic process), and GO:0006743 (ubiquinone metabolic process).

### Temperature and Salt-Dependent Growth Measurements

To assess how the growth of large-evolved (LS) and no selection (NS) populations responds to novel environmental conditions, growth was measured in M3 media across varying temperatures (27°C, 30°C, 35°C) and NaCl concentrations (0 M, 0.2 M, 0.4 M). Populations were derived from frozen stocks collected on day 38 of the evolution experiment. For each replicate, frozen stocks were grown to saturation in YPD, then inoculated into 6 mL of M3 media and incubated at 30°C for approximately 4-6 hours. Then cultures were diluted into fresh 6 mL M3 media at a standardized starting density of approximately 1.5 × 10⁶ cells. For salt conditions, NaCl was added to achieve final concentrations of 0.2 M and 0.4 M. Optical density at 600 nm (OD₆₀₀) was monitored over 48 hours at 10-minute intervals using a TVS062CA (Advantec MFS) compact rocking incubator set to 70 rpm at the specified temperature. Following 48 hours of incubation, 24 µL of each culture was transferred into 6 mL of fresh M3 medium to initiate the next growth period. This transfer was repeated after an additional 48 hours of incubation, at 96 hours total. Thus, each population was monitored across three consecutive 48-hour growth periods over a 6-day experiment. With two independently evolved populations represented for each selection regime, this resulted in six growth curves per selection regime across the 6-day period. For NaCl experiments, incubation was performed at 30°C; thus, the 0 M NaCl condition corresponds to the same environmental condition as 30°C in temperature experiments.

### Quantification of Growth Dynamics at the Transition from Exponential Phase

To quantify growth dynamics at the transition from exponential to stationary phase, the rate of change in OD₆₀₀ was calculated for each replicate using a sliding window of 110 minutes. The transition point was defined as the time corresponding to the minimum rate of change following the maximum rate of change. To avoid confounding effects from the stationary phase, where growth rates approach zero, upper time cutoffs were applied. For replicates in which the maximum rate of change occurred prior to 25 hours, analysis was restricted to the first 30 hours. For those in which the maximum occurred after 25 hours, a cutoff of 40 hours was applied.

These thresholds were determined based on visual inspection of growth trajectories to exclude the stationary phase while retaining biologically relevant transitions, including delayed growth associated with diauxic shifts. OD₆₀₀ values were then averaged across the 110-minute window centered on the identified transition time point for each replicate. These values were subsequently averaged across replicates within each condition for comparison between populations.

## Data Availability Statement

Data processing and analysis scripts, as well as processed alignment data can be found on the Open Science Framework: https://osf.io/2s3q7/overview?view_only=3ce9aad5ec5d4c1aaaca84654af5fc52. Raw fastq files for genomes can be accessed through NCBI SRA PRJNA1519969. All subsequent analyses performed in this publication can be reproduced directly with these data.

## Acknowledgments

Funding for this work was provided by a National Science Foundation Biological Integration Institution grant 2119963 (to KGS and ML). We would also like to thank the members of the ASU Biodesign Center for Mechanisms of Evolution and the Geiler-Samerotte Lab for their support and feedback.

## Supplemental Figures

**Figure S1:**
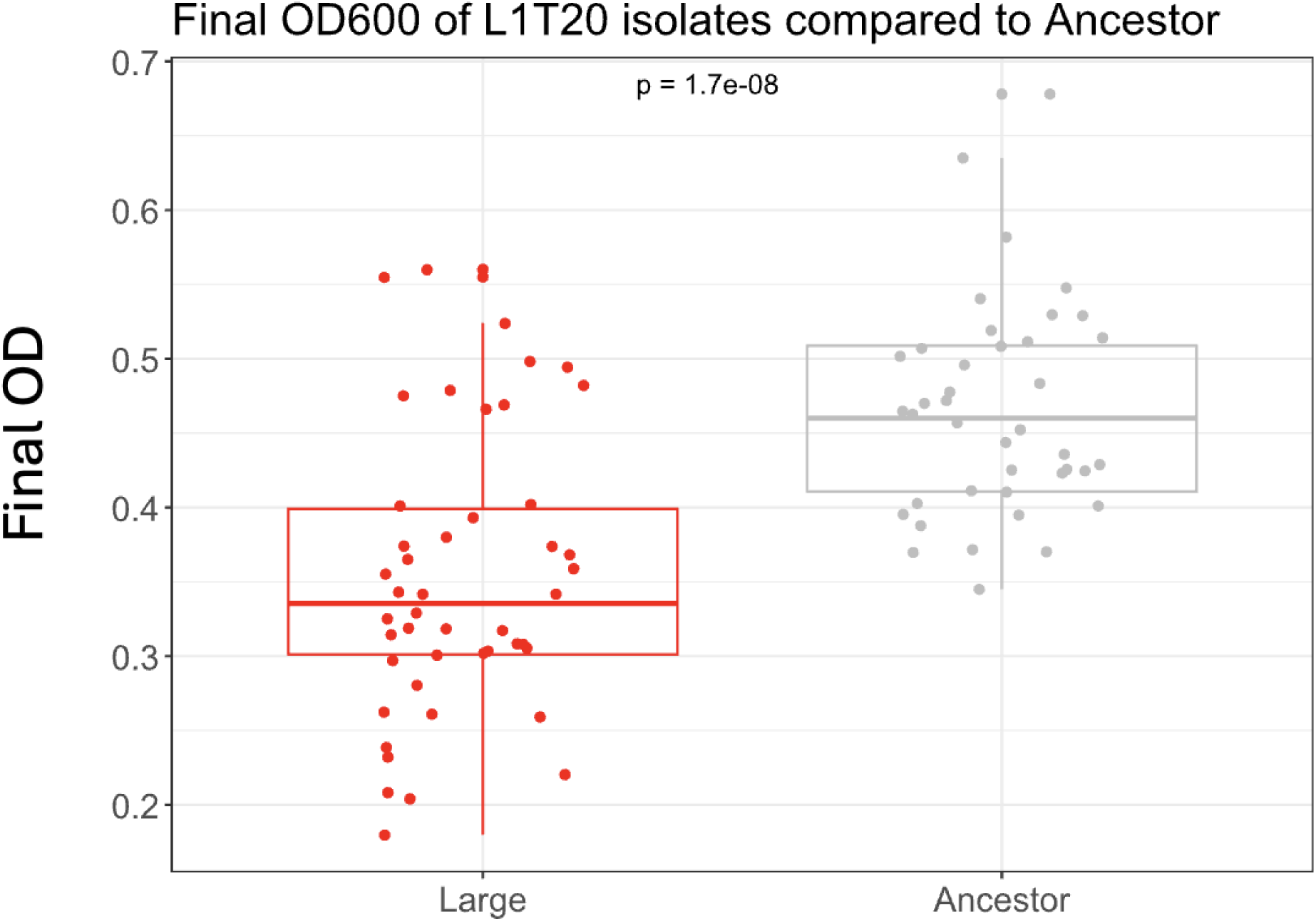
Evolved large isolates reach a lower final OD following 48 hours of growth. Every point is a cultured isolate initiated from a freezer stock where 20 uL was transferred into 200 uL of predefined M3 media. These isolates were clonally propagated in a 96 well plate and OD was measured after 48 hours. Final OD after 48 hours was measured for isolates from the evolved large population (n=46) and its ancestor population (n=40). Comparisons between Large and Ancestor isolates illustrate a statistically significant lower Final OD being reached in the Large isolates following 48 hours of growth.

**Figure S2:**
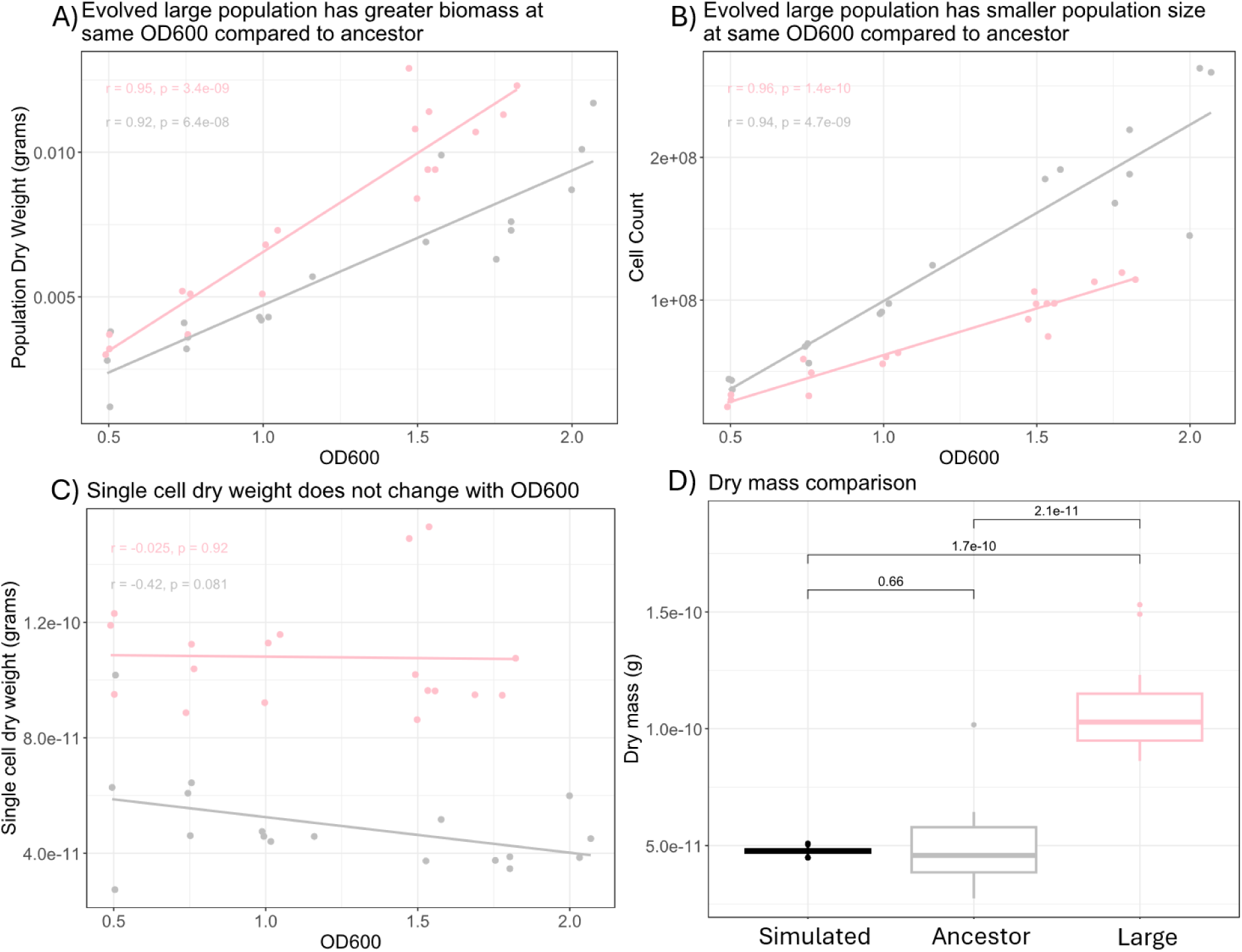
Evolved large populations reach a higher population biomass despite having a lower population size compared to the ancestor population. To determine how biomass production and the population size compared between the evolved large and its ancestral population, dry weight and cell count measurements were taken across different OD measurements during incubation. (A) Population dry weight measurements for the evolved large and its ancestor population in grams at different OD values. Linear regressions of both populations plotted with Pearson correlation coefficient plotted with their corresponding p values. Relationships show the large evolved population consistently reaching higher population biomass compared to the ancestor at approximately similar OD values. **(B)** Relationship between cell count and OD for both populations, with corresponding Pearson correlation coefficient and p values. Relationships show that ancestor population consistently reaches a higher cell count compared to large populations at approximately similar OD values. **(C)** Single cell dry weight measurements in grams across different OD values for both populations, with corresponding Pearson correlation coefficient and p values. Single cell dry weight was calculated by dividing the population dry weight by cell count. Relationships show no statistically significant change over different OD measurements with the large population consistently having higher single cell dry weight measurements compared to its ancestor at approximate OD values. **(D)** Comparisons between large and ancestor single cell dry mass measurements compared to simulated data of null expectation (Łabędź et al. 2017). Ancestor population shows no statistical difference compared to null expectation.

**Figure S3:**
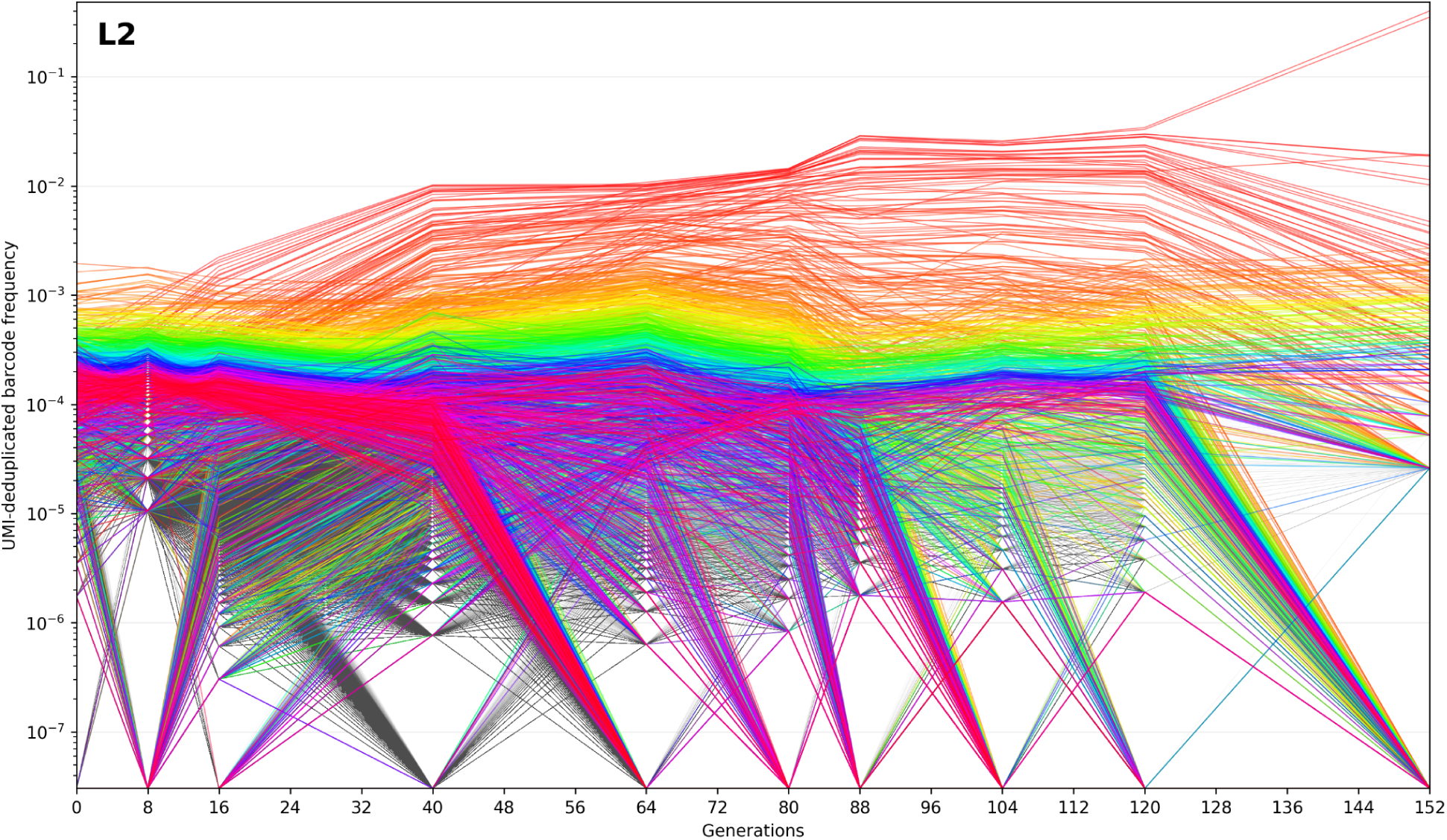
Barcode lineage dynamics across 152 generations in L2. For one replicate of our evolution experiment (large-size selected replicate 2 “L2”) we deeply sequenced the barcodes at many timepoints to understand their evolutionary dynamics. This graph shows UMI-deduplicated barcode frequencies across generations in L2. Each line represents one barcode lineage; the 2,000 most abundant lineages are colored and the remaining plotted lineages are gray. Low-depth time points (24, 32, and 48 generations) were excluded. In total, the 75,000 most abundant barcodes are shown.

**Figure S4:**
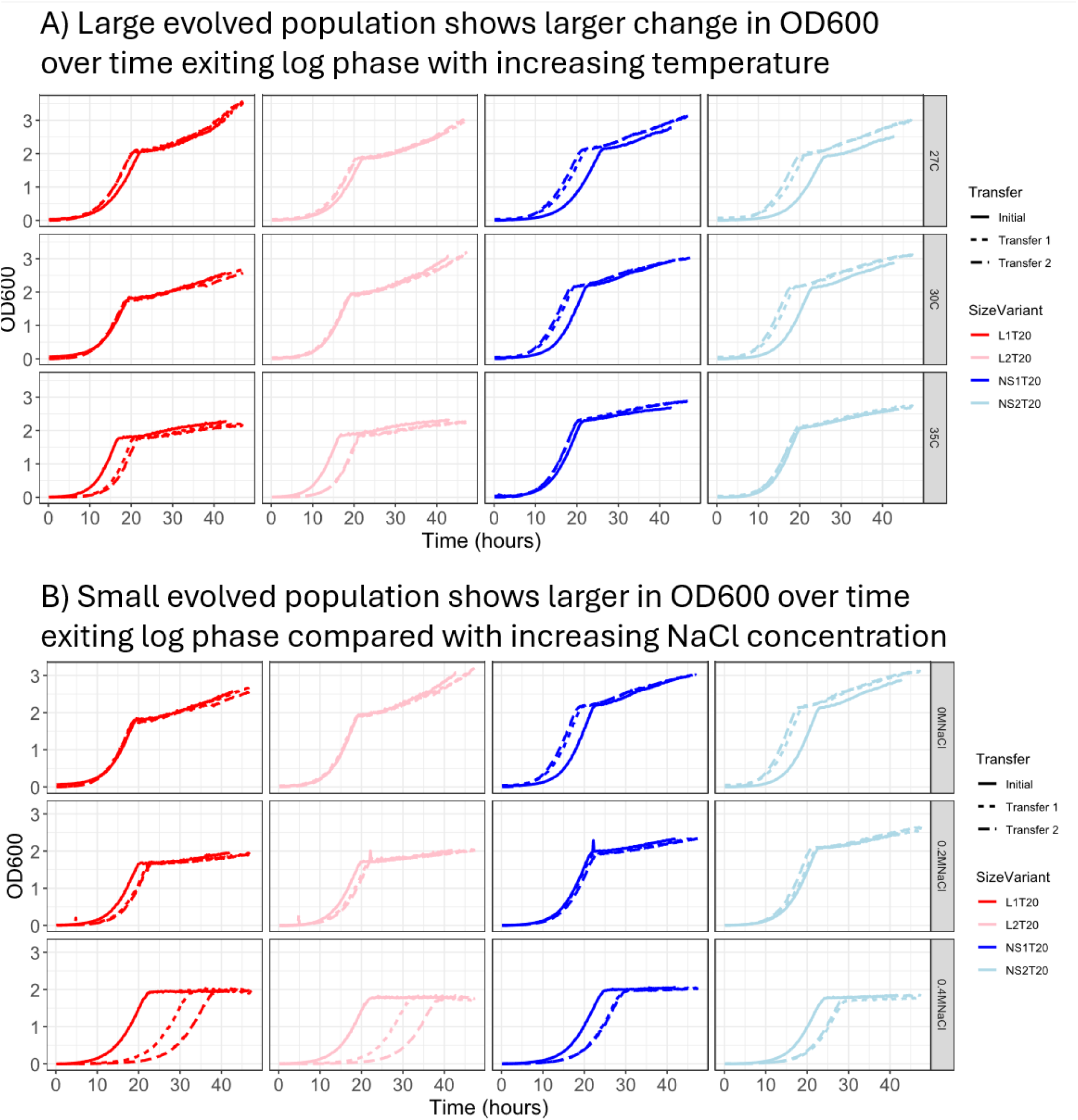
Raw growth curves of evolved populations across different temperature and salt environments. **(A)** Growth curves of each evolved population in 6 mL of M3 media at increasing temperatures, with 24 uL of culture transferred into fresh media every 48 hours twice. Large populations (red) at each transfer show lower OD values upon exiting exponential phase at increasing temperatures compared to the no selection populations (blue) at their respective transfers. **(B)** Growth curves of each evolved population in 6 mL of M3 media at increasing NaCl concentrations, with 24 uL of culture transferred into fresh media every 48 hours twice. NS populations show declining OD values at increasing salt concentrations while LS populations decline less strongly. All populations were sampled from frozen culture at day 38 of the evolution experiment.

